# Long-term Population Dynamics and Conservation of Proboscis Monkeys (*Nasalis larvatus*) in the Lower Kinabatangan, Sabah, Malaysia

**DOI:** 10.64898/2026.10.10.758256

**Authors:** Muhammad Nur Fitri-Suhaimi, Hiroshi Okamura, Nicola Karen Abram, Marc Ancrenaz, Benoit Goossens, Primus Lambut, Joseph Tangah, John C. M. Sha, Henry Bernard, Vijay Subbiah Kumar, Ikki Matsuda

## Abstract

Long-term ecological monitoring is essential for evaluating the effectiveness of conservation actions for long-lived tropical primates, yet few studies have integrated repeated population censuses with habitat-change analyses. We evaluated long-term population and habitat dynamics of the endangered proboscis monkey (*Nasalis larvatus*) in the Lower Kinabatangan, Sabah, Malaysian Borneo, by integrating a landscape-wide population survey conducted in 2025 with previous surveys from 2005 and 2014, long-term monitoring at three representative sites, and spatial analyses of land-use and land-cover change. Standardized late-afternoon boat-based surveys covering approximately 223 km of the Kinabatangan River and its tributaries recorded an estimated 2,420 individuals in 174 social groups, representing an increase of approximately 66% and 72% respectively compared with the 2005 survey. Group sizes (number of individuals per social group) remained largely unchanged despite the overall increase in population size. Long-term monitoring at three representative sites indicated stable population trajectories throughout the study period. Habitat analyses further revealed an increase in natural vegetation and limited conversion of forest to oil palm within protected areas between 2005 and 2025. Together, these independent lines of evidence indicate that the Lower Kinabatangan proboscis monkey population has recovered while maintaining overall demographic stability under long-term habitat protection. Nevertheless, substantial areas of suitable habitat remain outside the protected-area network, highlighting opportunities for additional habitat restoration and improved landscape connectivity. Our findings demonstrate the value of integrating long-term ecological monitoring with landscape-scale habitat assessment to inform adaptive conservation management in fragmented tropical landscapes.

## 1. Introduction

Evaluating the effectiveness of conservation actions requires more than documenting changes in population size; it also requires understanding how populations respond to long-term changes in habitat and landscape structure (Chapman and Peres, 2021; Geldmann et al., 2021; Le Saout et al., 2013). Long-term ecological monitoring provides one of the most powerful approaches for evaluating population trajectories, identifying the mechanisms underlying population change, and assessing the effectiveness of conservation interventions (Barelli et al., 2023; Chapman et al., 2017; Clutton-Brock and Sheldon, 2010). When integrated with spatial information on habitat change, such monitoring can reveal how wildlife populations respond to dynamic landscapes and provide an evidence-based foundation for adaptive conservation management (Chapman et al., 2010; Chapman and Peres, 2021). Despite its importance, long-term datasets that combine repeated population surveys with habitat assessments remain rare for tropical wildlife, particularly long-lived primates, because they require sustained investment over many years (Atmoko et al., 2024; Bernard et al., 2025b; Matsuda et al., 2020; Mohd-Daut et al., 2021). Although several long-term monitoring programs have been established for proboscis monkeys in Sabah, including the Klias Peninsula (Bernard et al., 2025b, 2021), few studies have integrated repeated landscape-wide population censuses with habitat-change analyses to evaluate long-term population and habitat dynamics across an entire riverine landscape.

The proboscis monkey (*Nasalis larvatus*) is an endangered, highly social colobine endemic to Borneo that is strongly associated with riverine and other wetland forests (Boonratana et al., 2021; Meijaard and Nijman, 2000). The species typically forms one-male– multi-female groups and all-male groups, although multi-male–multi-female groups and solitary males are occasionally observed (Boonratana, 2002; Matsuda et al., 2020; Murai, 2004). As predominantly folivorous primates, proboscis monkeys rely on a variety of wetland forest resources, including young leaves, shoots, fruits, seeds, and flowers (Bernard et al., 2019; Matsuda et al., 2009a; Yeager, 1989). A distinctive ecological characteristic of the species is its strong dependence on riverine habitats. Individuals typically forage up to approximately 800– 1,000 m inland during the day before returning to riverside sleeping trees each evening, where multiple social groups may temporarily aggregate into sleeping bands (Bernard et al., 2011; Matsuda et al., 2024, 2010, 2009b). This predictable use of riparian habitats enables standardized boat-based surveys and makes the species particularly well suited for long-term population monitoring at the landscape scale.

Despite legal protection throughout its range, the proboscis monkey continues to face substantial conservation challenges. The species has been listed as Endangered on the IUCN Red List since 2000 and is protected under Appendix I of the Convention on International Trade in Endangered Species (CITES), while in Sabah it is classified as a Totally Protected Species under the Sabah Wildlife Conservation Enactment (Boonratana et al., 2021; Sabah Wildlife Conservation Enactment, 1997; UNEP-WCMC, 2003). Nevertheless, habitat loss, fragmentation, and degradation remain the principal threats to its long-term persistence, driven primarily by the conversion of lowland wetland forests for agriculture, particularly oil palm plantations, together with aquaculture and urban development (Atmoko et al., 2024; Bernard et al., 2025b; Bernard and Amidy, 2025; Gaveau et al., 2014; Matsuda et al., 2020; Toulec et al., 2020). Because the species depends heavily on riparian habitats, these landscape changes can disproportionately affect its distribution and population dynamics (Matsuda et al., 2010, 2009b). Consequently, evaluating long-term population trends together with habitat change is essential for determining whether existing conservation measures are achieving their intended outcomes and for identifying priorities for future management (Bernard et al., 2025a, 2025b, 2021; Chapman and Peres, 2021).

Recognizing both the urgent conservation needs of the species and its socio-economic importance, particularly its value for ecotourism and local livelihoods (Leasor and Macgregor, 2014), the Sabah State Government developed the Proboscis Monkey Action Plan for Sabah (2019–2028) (Sabah Wildlife Department, 2019). The plan identifies seven priority populations for long-term conservation, including the Lower Kinabatangan, and emphasizes two complementary objectives: protecting remaining habitat and establishing systematic long-term population monitoring. Consequently, evaluating long-term population and habitat trends within these priority landscapes is essential for assessing the effectiveness of current conservation actions and informing future management decisions. However, such integrated long-term studies remain scarce because they require sustained logistical commitment and substantial investments of time and resources (Atmoko et al., 2024; Bernard et al., 2025a, 2021; Matsuda et al., 2020; Toulec et al., 2022). This knowledge gap is particularly acute in tropical floodplain ecosystems, where rapid landscape change requires repeated assessments of both population status and habitat condition to support adaptive, evidence-based conservation management.

Landscape-wide surveys conducted in the Lower Kinabatangan in 2005 and 2014 established baseline information on the abundance and distribution of proboscis monkeys, identifying the region as one of the most important strongholds for the species in Sabah (Matsuda et al., 2020; Sha et al., 2008). These surveys indicated a period of relative population stability accompanied by limited habitat loss (Matsuda et al., 2020). However, no comprehensive reassessment has been undertaken for more than a decade. Given the rapid pace of landscape change in tropical floodplain ecosystems, it remains unknown whether this apparent stability has been sustained and whether existing conservation measures have remained effective.

To address this knowledge gap, we conducted a comprehensive landscape-wide survey in 2025 and integrated the results with long-term monitoring data and habitat-change analyses. Specifically, our objectives were to: (1) evaluate changes in population size, distribution, and group structure by comparing the 2025 survey with previous surveys conducted in 2005 and 2014; (2) examine long-term population trends at three representative monitoring sites; and (3) quantify land-use and land-cover changes between 2005 and 2025 to evaluate habitat dynamics and identify priorities for future landscape conservation.

## 2. Material and methods

### 2.1. Study area

The Lower Kinabatangan floodplain in eastern Sabah, Malaysian Borneo (5°10′–5°50′N; 117°40′–118°30′E), comprises a mosaic of riverine, freshwater swamp, mangrove, and nipa palm forests embedded within a heavily modified tropical floodplain landscape dominated by oil palm plantations and selectively logged forests (Abram et al., 2014; Ancrenaz et al., 2004). Since the early 2000s, approximately 27,000 ha of fragmented forest along the Kinabatangan River has been protected under the Lower Kinabatangan Wildlife Sanctuary, which comprises 10 forest blocks embedded within a landscape containing additional protected and unprotected forest patches with varying levels of degradation (Abram et al., 2014; Ancrenaz et al., 2004).

### 2.2. Survey of Kinabatangan River and tributaries

Proboscis monkey surveys followed the standardized boat-based protocol described by Matsuda et al. (2020), ensuring methodological consistency with surveys conducted in 2005 and 2014. Surveys were conducted exclusively during the late afternoon, when proboscis monkeys aggregate at riverside sleeping sites and population estimates are most reliable (Matsuda et al., 2016).

We conducted 21 boat surveys over 18 field days between 14 September and 1 October 2025, covering 222.72 km of the Kinabatangan River and its tributaries (Figure 1). The mean distance covered per boat survey was 13.4 ± 3.7 km. Survey effort increased from 186.2 km in 2014 to 222.72 km in 2025, with the additional 36.52 km. The additional 36.52 km consisted primarily of mangrove sections that had not been surveyed previously and therefore increased the spatial coverage of the census without altering the core survey transects established in earlier surveys. Surveys were conducted from 16:30 until sunset, and each river section was surveyed only once (Matsuda et al., 2020; Sha et al., 2008). Adjacent river sections were surveyed on the same or consecutive days to minimize the possibility of duplicate counts resulting from movements of social group between neighbouring river sections.

**Fig. 1.**
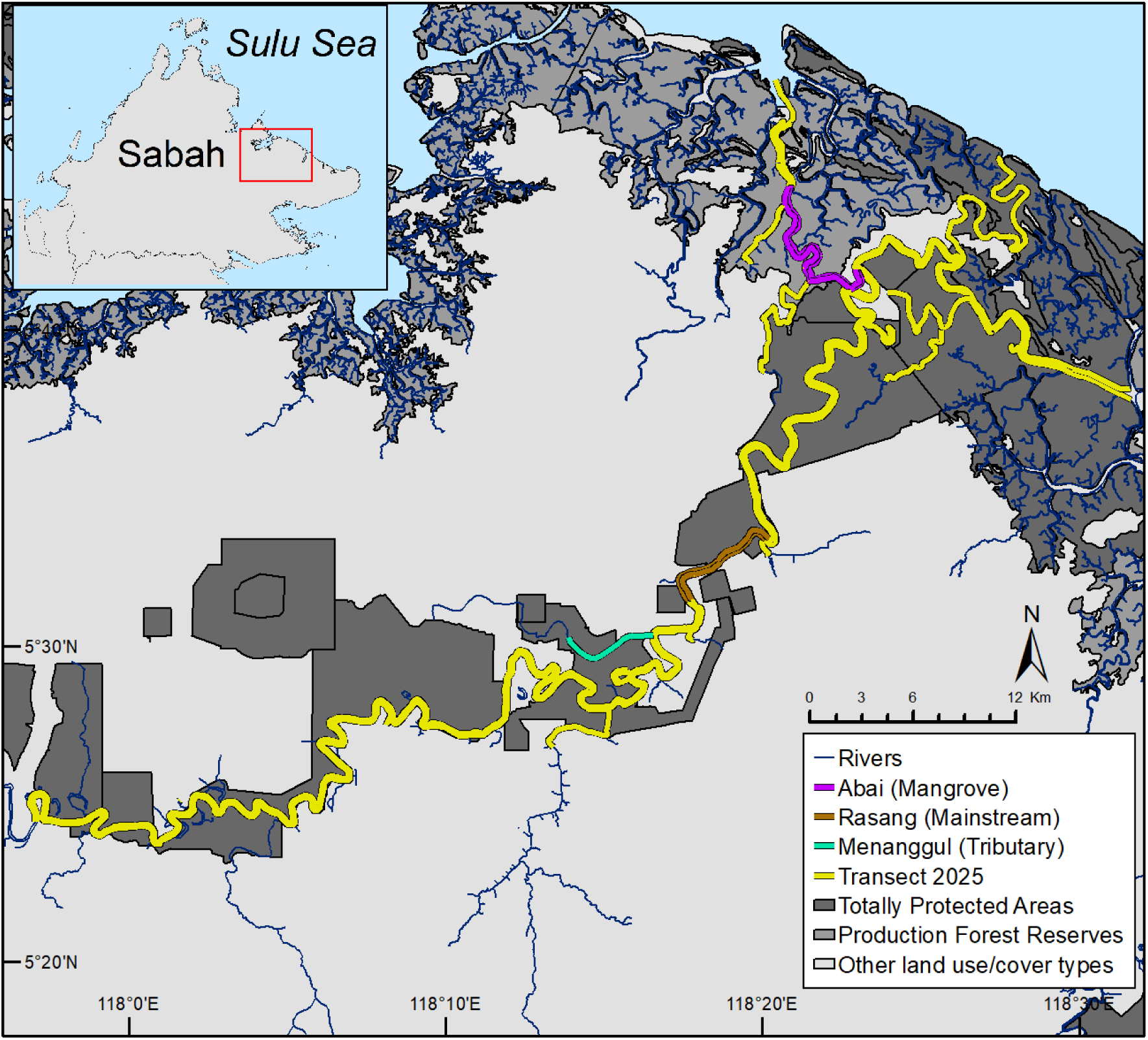
Map of the Lower Kinabatangan (eastern Sabah, Malaysian Borneo), showing the 2025 survey transect and the locations of the three long-term monitoring sites.

For each sighting, we recorded group size and composition, classifying groups as one-male–multi-female groups (OMG), all-male groups (AMG), multi-male–multi-female groups, or solitary individuals (Bennett and Sebastian, 1988; Murai, 2004). Individuals whose age-sex class could not be reliably determined were classified as undetermined. Survey routes and sighting locations were recorded using a handheld GPS.

### 2.3. Long-term monitoring sites

In addition to the landscape-wide survey, late-afternoon boat surveys were conducted at three long-term monitoring sites representing the major habitat types in the Lower Kinabatangan (Figure 1). Menanggul, a 6.0-km riverine tributary surveyed 945 times between 2005 and 2025. Survey covered the full 6.0-km section until 2014, and the lower 4.0-km stretch thereafter. Rasang, a 13.2-km section of the main river surveyed 352 times between 2007 and 2023. Abai, a 13.1-km mangrove section surveyed 285 times between 2010 and 2025. All analyses were restricted to surveys conducted between 16:00 and 18:00 to control for time-of-day effects on detectability.

### 2.4. Statistical analysis

#### 2.4.1. Population trends

We compared the numbers of individuals and groups recorded in the 2025 survey with those reported in 2005 (Sha et al., 2008) and 2014 (Matsuda et al., 2020). Differences among survey years were evaluated using chi-squared goodness-of-fit test (Zar, 1999), with the 2005 survey serving as the reference.

#### 2.4.2. Group size

Changes in group sizes were analysed separately for OMGs and AMGs using generalized linear models with a quasi-Poisson error distribution and log link function. Group size was fitted as the response variable. Survey year (2005, 2014, and 2025) was included as a fixed-effect, with 2005 specified as the reference. Overdispersion was assessed using a year-specific dispersion parameter (Kleiber and Zeileis, 2008).

#### 2.4.3. Long-term monitoring sites

Long-term population trends at the three monitoring sites were analysed using a state-space model that separates observation error from stochastic population process (Buckland et al., 2004). Observed counts were modelled using a Poisson observation model linked to a first-order random walk process model. Model parameters were estimated by a maximum likelihood method using the R package Template Model Builder (Kristensen et al., 2016). All statistical analyses were conducted in R version 4.4.1. The full model specification is provided in the Supplementary Methods (S1).

### 2.5. Temporal land use and land cover within potential proboscis monkey habitat

To quantify land-use and land-cover change between 2005 and 2025, we used the 2005 and 2014 land-cover datasets developed by Matsuda et al. (2020). The 2025 land-cover layer was generated by updating the 2014 dataset using cloud-free Sentinel-2 satellite imagery acquired in 2024–2025. The 2014 and 2025 land-cover layers were derived from higher-resolution satellite imagery than the 2005 layer, allowing a more detailed and accurate classification of land-cover types. Because survey length differed slightly across years, and to keep the land-use change analyses consistent with Matsuda et al. (2020), we used the same transect coverage and 800-m buffer around the riverbank (totalling 25,795 ha) to identify any habitat change. Within this area, 15 land-cover classes were grouped into three major categories: forest/natural vegetation, oil palm plantations, and settlements. Each land-cover category was further classified as either protected (Lower Kinabatangan Wildlife Sanctuary and forest reserves) or unprotected (Abram et al., 2014).

## 3. Results

### 3.1. Landscape-wide population status

We recorded 2,420 proboscis monkeys in 174 groups across the Lower Kinabatangan in 2025. These comprised 119 OMGs, 46 AMGs, including solitary males, eight multi-male–multi-female groups, and one group of undetermined composition. Of the total individuals recorded, 262 were recorded in mangrove areas (including beach forest, mangrove forest, nipa palm forest, and mixed mangrove–nipa forests; hereafter referred to as “mangrove forests”), and 2,158 were observed in riverine areas (including freshwater swamp forest, swamp, peat swamp forest, and lowland dry forest; hereafter referred to as “riverine forests”).

Mean group size was 15.4 ± 5.0 individuals (range = 2–25) for OMGs and 9.0 ± 4.5 individuals (range = 1–16) for AMGs (Table 1). OMGs were significantly smaller in mangrove forests (10.6 ± 4.7, n = 19) than in riverine forests (16.4 ± 4.5, n = 100; Mann–Whitney U test: W = 353, *p* < 0.001). In contrast, AMG size did not differ significantly between mangrove (7.8 ± 3.8, n = 4) and riverine forests (9.1 ± 4.5, n = 42; W = 71.5, *p* = 0.64).

**Table 1.**
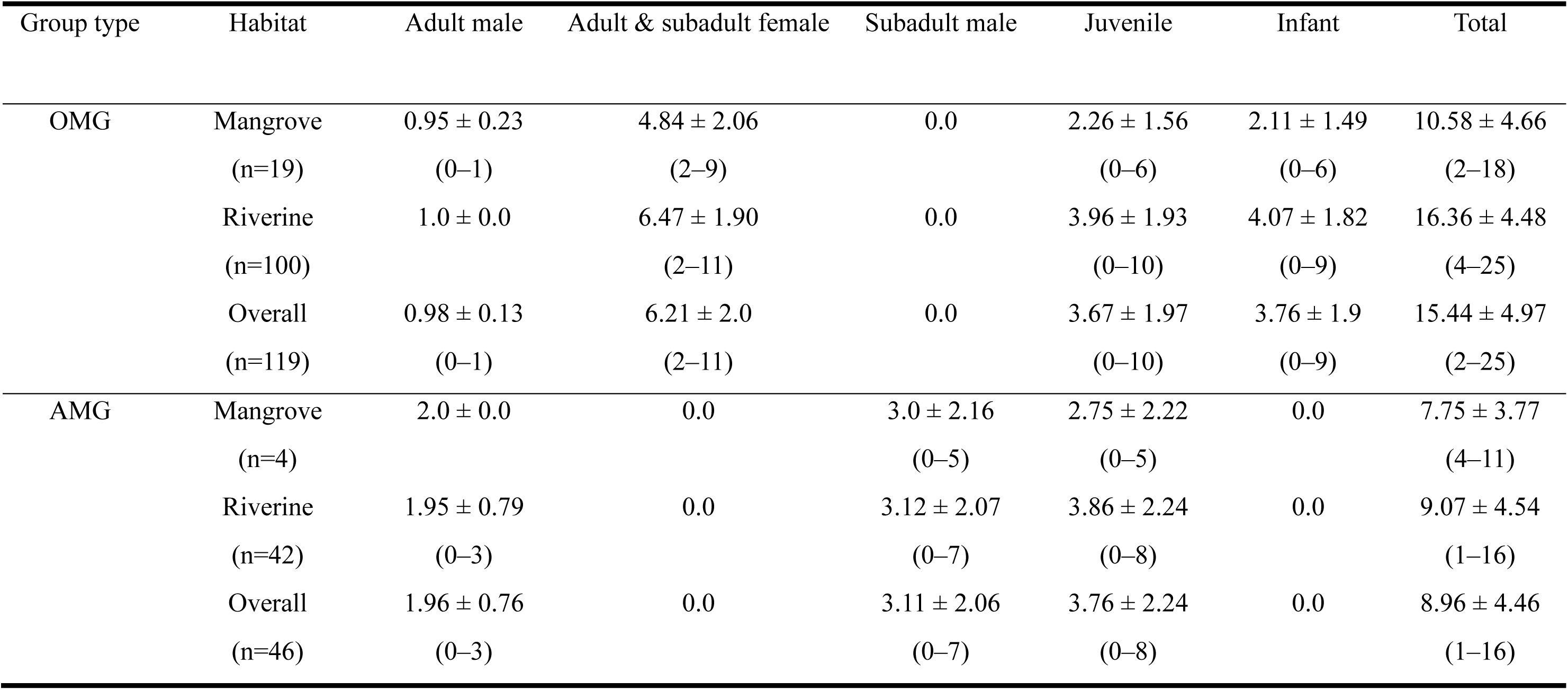
Composition and group size (mean ± SD, range) of one-male–multi-female groups (OMG) and all-male groups (AMG) recorded during the 2025 survey in the Lower Kinabatangan.

Estimated population size increased from 1,454 individuals in 2005 (Sha et al., 2008) to 1,980 in 2014 (Matsuda et al., 2020) and 2,420 in 2025 (this study; *χ*² = 240.1, *df* = 2, *p* < 0.001). Similarly, the number of observed groups increased from 101 in 2005 to 164 in 2014 and 174 in 2025 (*χ*² = 22.8, *df* = 2, *p* < 0.001).

Poisson regression analyses showed that mean OMG size was significantly smaller in 2014 than in 2005 (*p* = 0.0045), but did not differ between 2005 and 2025 (*p* = 0.81; Table 2, Figure 2). No significant differences in AMG size were detected among survey years.

**Fig. 2.**
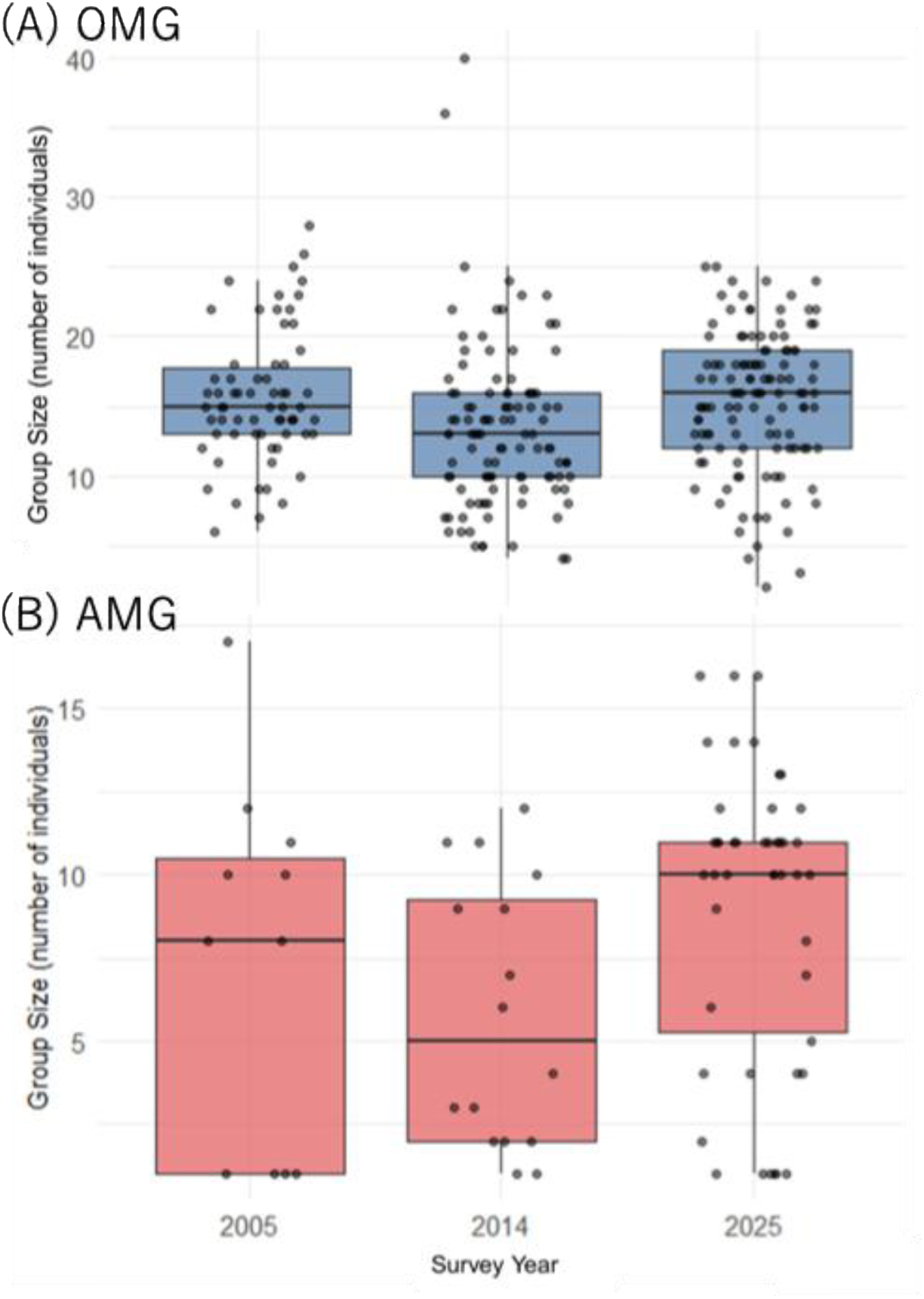
Changes in group size of one-male–multifemale groups (A) and all-male groups (B) across the 2005, 2014, and 2025 surveys. Boxes represent the interquartile range (25th–75th percentiles), horizontal lines indicate the median, whiskers extend to the most extreme values within 1.5 × the interquartile range, and points represent individual group sizes.

**Table 2.**
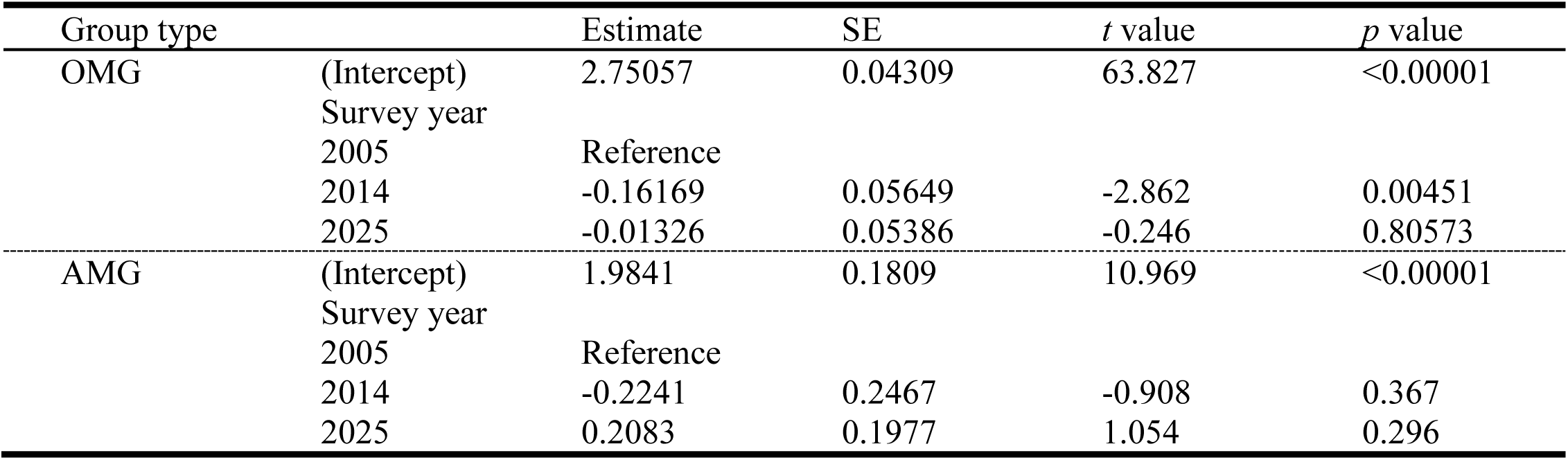
Generalized linear model results for changes in the size of one-male–multi-female groups (OMG) and all-male groups (AMG) across the 2005, 2014, and 2025 surveys.

### 3.2. Long-term population dynamics

State-space models indicated no significant long-term population change at any of the three monitoring sites (Table 3, Figure 3). Estimated population trends were similar among Menanggul, Rasang, and Abai, with 95% confidence intervals overlapping zero.

**Fig. 3.**
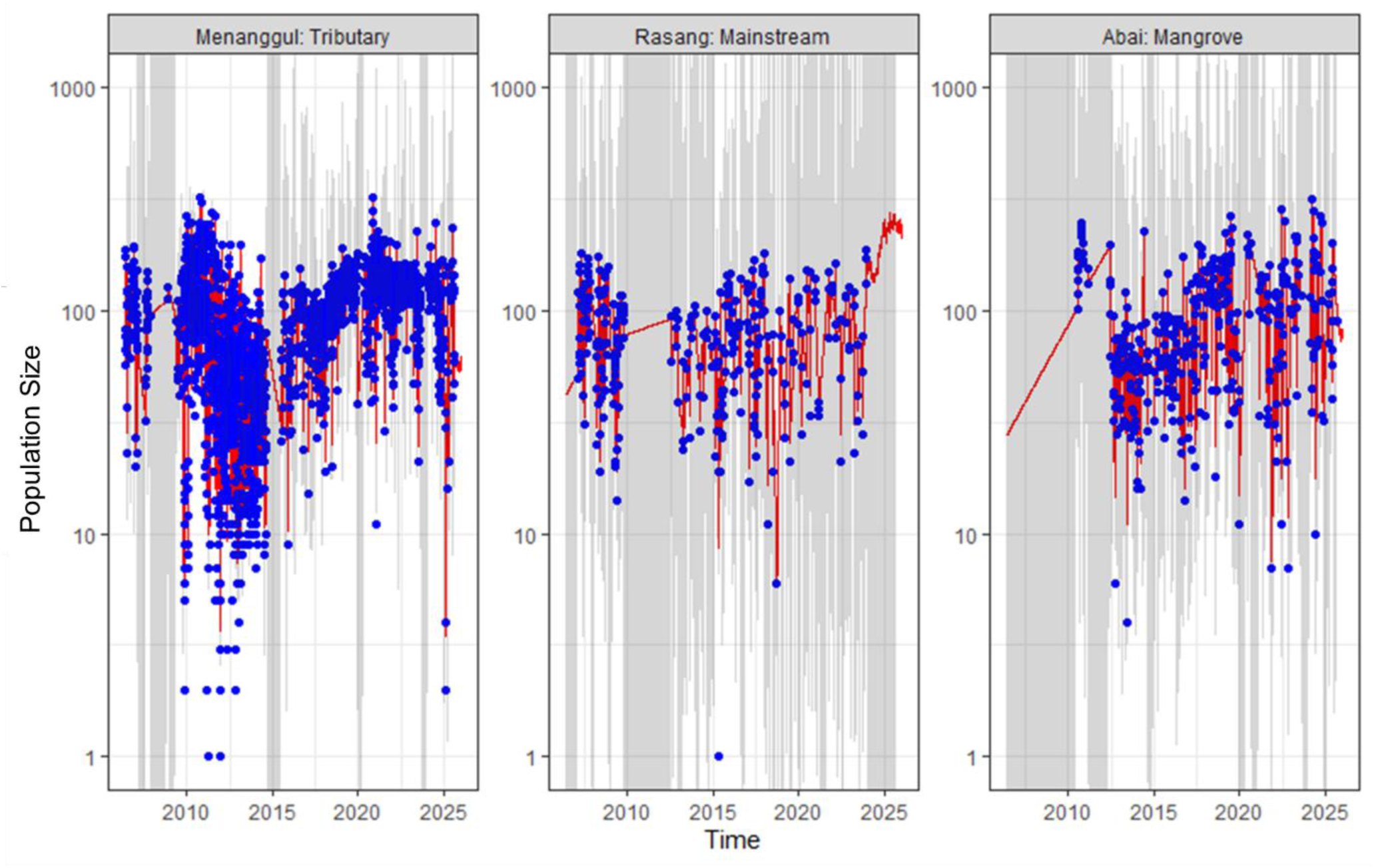
Long-term population dynamics of proboscis monkeys at three monitoring sites in the Lower Kinabatangan. Blue points represent observed counts, red lines represent state-space model estimates, and grey shaded areas indicate 95% confidence intervals. Panels show Menanggul (tributary), Rasang (mainstream), and Abai (mangrove).

**Table 3.** State-space model estimates of long-term population trends for the three long-term monitoring sites in the Lower Kinabatangan. Values are presented as trend estimates (μ), standard errors (SE), and 95% confidence intervals.

|  | Estimate | SE | 95% CI |
| --- | --- | --- | --- |
| Log sigma | -0.72602267 | 0.016692373 |  |
| Monitoring site mean |  |  |  |
| Trend estimate ($\mu$): Menanggul (Tributary) | 0.0005559291 | 0.003503583 | [-0.006311, 0.007423] |
| Trend estimate ($\mu$): Rasang (Mainstream) | 0.0005559423 | 0.003536100 | [-0.006375, 0.007487] |
| Trend estimate ($\mu$): Abai (Mangrove) | 0.0005559400 | 0.003503617 | [-0.006311, 0.007423] |

### 3.3. Habitat change within the potential proboscis monkey habitat

The potential habitat of proboscis monkeys, defined as an 800-m buffer from the riverbank, covered 25,795 ha in the Lower Kinabatangan (Figure 4). Natural vegetation accounted for 20,536 ha (79.6%) of this area, followed by 5,056 ha (19.6%) of oil palm plantations and 203 ha (< 1%) of settlements (Table 4). Overall, between 2005 and 2025, natural vegetation cover increased by 94 ha, oil palm cover decreased by 60 ha, and settlement cover increased by 203 ha; settlements concentrated primarily outside protected areas. Although these changes were marginal, they reflect a net gain in forest vegetation and a net reduction in oil palm cover.

**Fig. 4.**
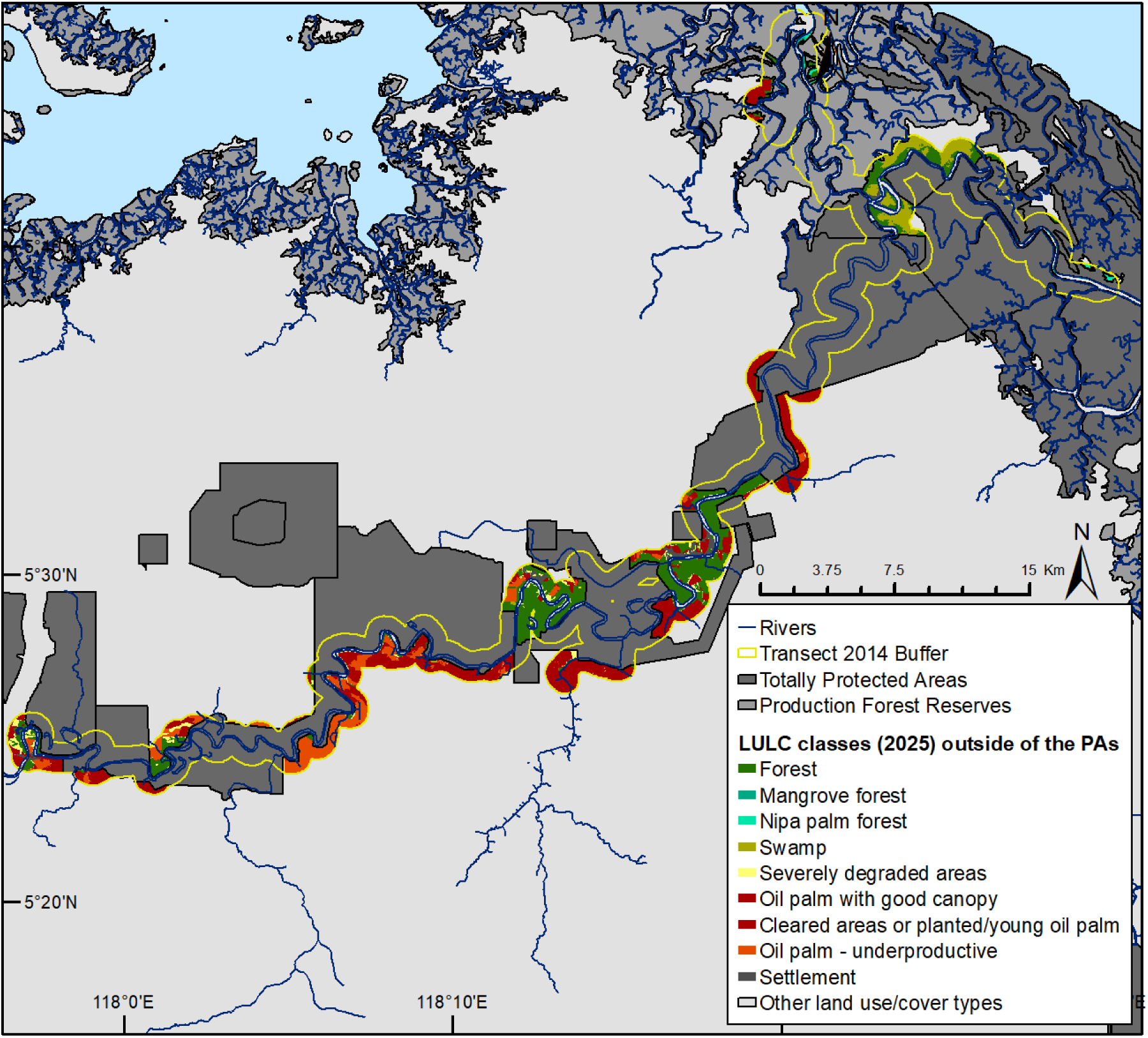
Land-use and land-cover within the potential habitat of proboscis monkeys in the Lower Kinabatangan in 2025. Protected areas are outlined in dark grey, illustrating the distribution of unprotected natural vegetation and oil palm plantations.

**Table 4.** Land-use and land-cover within the potential habitat of proboscis monkeys (800-m buffer) in 2005, 2014, and 2025.

| Land-use/land-cover type within potential proboscis monkey range | Total extent in 2005 in ha (%) | Total extent in 2014 in ha (%) | Total extent in 2025 in ha (%) | Difference from 2025 to 2005 |
| --- | --- | --- | --- | --- |
| **Overall** |  |  |  |  |
| Forest/natural vegetation | 20,442 (79.9%) | 20,591 (79.8%) | 20,536 (79.6%) | 94 |
| Oil palm | 5,116 (20.0%) | 5,018 (19.5%) | 5,056 (19.6%) | -60 |
| Settlement | 0 | 189 (0.73%) | 203 (0.79%) | 203 |
| **Total** | **25,558** | **25,798** | **25,795** | **-** |
| **Protected area** |  |  |  |  |
| Forest/natural vegetation | 15,878 (97.5%) | 16,136 (98.0%) | 16,145 (98.0%) | 267 |
| Oil palm | 414 (2.50%) | 335 (2.00%) | 326 (1.98%) | -88 |
| Settlement | 0 | 0 | 1 (<0.1%) | 1 |
| **Total** | **16,292** | **16,471** | **16,472** | **-** |
| **Outside of protected area** |  |  |  |  |
| Forest/natural vegetation | 4,564 (49.3%) | 4,455 (47.8%) | 4,391 (47.1%) | -173 |
| Oil palm | 4,702 (50.7%) | 4,683 (50.2%) | 4,730 (50.7%) | 28 |
| Settlement | 0 | 189 (2.03%) | 202 (2.17%) | 202 |
| **Total** | **9,266** | **9,327** | **9,323** | **-** |

Protected areas encompassed 16,472 ha of the potential habitat, leaving the remaining 9,323 ha outside protected areas. Within these areas, natural vegetation increased by 267 ha between 2005 and 2025, whereas oil palm plantations declined by 88 ha (Table 4). In contrast, natural vegetation outside protected areas declined by 173 ha over the same period, accompanied by increases in oil palm plantations (+28 ha) and settlements (+202 ha). Notably, the potential habitat of proboscis monkeys spans both protected and outside protected areas.

Outside protected areas, natural vegetation cover declined substantially between 2005 and 2014, from 4,564 ha to 4,455 ha; this represents a net loss of 109 ha (Table 5). However, from 2014 to 2025, the natural vegetation cover remained relatively stable, decreasing only marginally from 47.8% to 47.1% of the total outside protected areas. Despite the limited forest loss recorded between 2014 and 2025, 4,391 ha of natural vegetation within the potential habitat remained outside protected areas (Table 5). This included 70 ha of mangrove forest, 204 ha of nipa palm, and 604 ha of swamp habitat, all of which represent important habitats for proboscis monkeys and remain relatively unchanged from 2014 to 2025, despite being outside protected areas. In addition, 326 ha of oil palm plantations remained within protected areas in 2025, of which 93 ha were classified as underproductive because they occurred on seasonally inundated land unsuitable for oil palm cultivation (Abram et al., 2014). Outside protected areas, a further 1,331 ha of underproductive oil palm plantations were identified on seasonally inundated land (Table 5).

**Table 5.** Distribution of land-use and land-cover types inside and outside protected areas within the potential habitat of proboscis monkeys in 2025, 2014 and 2025.

| Land-use/land-cover type within potential proboscis monkey range |  | Within the protected areas (ha) |  |  | Outside of the protected areas (ha) |  |  |
| --- | --- | --- | --- | --- | --- | --- | --- |
| Year |  | 2005 | 2014 | 2025 | 2005 | 2014 | 2025 |
| Forest/natural vegetation | Beach forest | - | 82 | 82 | - | 21 | 21 |
|  | Mangrove forest | - | 168 | 168 | - | 70 | 70 |
|  | Nipa palm forest | - | 2,301 | 2,301 | - | 204 | 204 |
|  | Transitional forest | - | 3,054 | 3,054 | - | 566 | 560 |
|  | Freshwater swamp forest | - | 5,210 | 5,210 | - | 997 | 974 |
|  | Seasonal freshwater swamp forest | - | 3,273 | 3,276 | - | 1,048 | 1,038 |
|  | Peat swamp forest | - | 43 | 43 | - | 9 | 9 |
|  | Swamp | - | 772 | 772 | - | 605 | 604 |
|  | Limestone forest | - | 70 | 70 | - | 76 | 75 |
|  | Lowland dry forest | - | 889 | 889 | - | 583 | 572 |
|  | Severely degraded areas (mixed forest types) | - | 274 | 280 | - | 276 | 264 |
|  | **Total** | **15,878** | **16,136** | **16,145** | **4,564** | **4,455** | **4,391** |
| Oil palm | Cleared areas or planted/young oil palm | - | 13 | 14 | - | 229 | 264 |
|  | Oil palm with good canopy | - | 227 | 219 | - | 3,134 | 3,135 |
|  | Oil palm (underproductive <25%) | - | 95 | 93 | - | 1,320 | 1,331 |
|  | **Total** | **414** | **335** | **326** | **4,702** | **4,683** | **4,730** |
| Settlement | Settlement | 0 | 0 | 1 | 0 | 189 | 202 |
|  | **Total** | **0** | **0** | **1** | **0** | **189** | **202** |

## 4. Discussion

### 4.1 Long-term population recovery under landscape-scale protection

Protected areas are widely recognized as the cornerstone of biodiversity conservation, yet relatively few studies have evaluated their long-term effectiveness using consistent ecological monitoring, particularly for long-lived tropical primates (Barelli et al., 2023; Chapman et al., 2017; Chen et al., 2023; Geldmann et al., 2021; Le Saout et al., 2013). Our study provides one of the longest integrated landscape-scale assessments of a tropical primate population, combining repeated population surveys, long-term monitoring at three representative sites, and habitat-change analyses throughout the study period. Together, these independent lines of evidence indicate that the proboscis monkey population in the Lower Kinabatangan has increased while maintaining overall demographic stability under long-term habitat protection.

Landscape-wide surveys showed that the estimated proboscis monkey population increased from 1,454 individuals in 2005, to 1,980 individuals in 2014 and 2,420 individuals in 2025, representing an increase of approximately 66% over two decades. Because all three surveys were conducted using the same standardized boat-based census protocol (Matsuda et al., 2020; Sha et al., 2008), this increase is unlikely to reflect methodological differences and instead provides strong evidence for a genuine long-term population recovery. Nevertheless, as with any long-term wildlife monitoring programme, changes in detection probability associated with observer teams or animal behaviour cannot be entirely excluded. It is also noteworthy that the additional 36.52 km surveyed in 2025 consisted predominantly of mangrove habitat and yielded only two additional social groups, indicating that the expanded survey coverage had only a minimal influence on the overall population estimate. Although boat-based surveys inevitably provide minimum population estimates, the consistency of survey design across all three census periods, together with the independent population trajectories observed at the three long-term monitoring sites, supports the conclusion that the observed increase reflects a genuine biological trend rather than methodological artefacts. This pattern contrasts with the relatively stable population trajectories reported from the Klias Peninsula and Balikpapan Bay, where long-term surveys have generally indicated limited net population change despite continued habitat pressures (Bernard et al., 2025b, 2021; Toulec et al., 2022). Importantly, this pattern is consistent with the long-term trends observed independently at the three monitoring sites, suggesting that the landscape-wide increase does not simply reflect localized shifts in distribution.

Despite the marked increase in overall population size, changes in group size were modest. Mean OMG size was significantly smaller in 2014 than in 2005, but this difference was no longer evident in the 2025 survey, suggesting that the temporary reduction observed a decade earlier may have been reversed. Likewise, AMG size remained relatively constant throughout the study period. Together with the stable population trajectories estimated independently from the three long-term monitoring sites, these findings suggest that population recovery was accompanied by the maintenance of normal social organization rather than by major changes in group structure. Notably, the increase in overall abundance appears to have been driven largely by an increase in the number of social groups rather than by marked increases in group size. Although boat-based surveys provide only a snapshot of social organization, the consistency of group size across surveys supports this interpretation. Our findings further emphasize the value of long-term ecological monitoring for detecting population responses in long-lived primates that may not be evident over shorter time scales (Chapman et al., 2017, 2010).

### 4.2 Protected areas are effective, but not sufficient

The habitat analyses provide strong evidence that long-term habitat protection has contributed substantially to the persistence and recovery of the Lower Kinabatangan proboscis monkey population. Within the 800-m buffer representing the species’ potential habitat, natural vegetation within protected areas increased by 267 ha between 2005 and 2025, while oil palm plantations declined by 88 ha. Although these changes represent only a small proportion of the total habitat, they indicate that suitable habitats for proboscis monkeys, including mangrove, nipa palm, and swamp forests, have been largely maintained within the protected-area network over the past two decades. These findings support the growing consensus that well-managed protected areas can effectively reduce habitat loss and promote biodiversity persistence, particularly when protection is maintained over sufficiently long time periods (Chapman and Peres, 2021; Geldmann et al., 2021; Le Saout et al., 2013). However, our results also demonstrate that protected areas alone are insufficient. Despite the overall positive trends, nearly 4,400 ha of suitable habitat within the potential habitat of proboscis monkeys remains outside the protected-area network, highlighting the importance of conservation approaches that extend beyond reserve boundaries and maintain suitable habitat across the wider landscape. Although forest loss within the 800-m buffer outside protected areas was substantial between 2005 and 2014, it remained relatively limited between 2014 to 2025. Our habitat analysis nevertheless identified several priorities for future landscape-scale conservation. A total of 4,391 ha of suitable habitat remains outside the current protected-area network, including mangrove, nipa palm, and swamp forests that provide important feeding, sleeping, and ranging habitats for proboscis monkeys while also contributing to habitat connectivity across the fragmented floodplain (Bennett and Sebastian, 1988; Bernard et al., 2019; Matsuda et al., 2009a). In addition, we identified 1,331 ha of underproductive oil palm plantations on seasonally inundated land outside protected areas, together with a further 93 ha within protected areas. Because these areas are poorly suited for commercial oil palm cultivation, they represent realistic opportunities for habitat restoration or strategic expansion of habitat protection with relatively limited conflict with agricultural production. Increasingly, conservation planning has emphasized that long-term biodiversity persistence depends not only on protected areas but also on maintaining habitat connectivity and restoring priority habitats throughout the surrounding landscape matrix (Gaveau et al., 2014; Hearn et al., 2019). Our findings provide spatially explicit evidence to prioritize future restoration efforts by identifying both unprotected suitable habitats and underproductive oil palm plantations that represent realistic opportunities for habitat restoration and landscape-scale conservation. These results provide a practical foundation for adaptive conservation planning that complements the existing protected-area network while promoting long-term conservation across the wider Lower Kinabatangan landscape (Abram et al., 2014).

### 4.3 Conservation implications for riparian landscapes

The present study further highlights the importance of conserving riparian habitats and other suitable habitats that support proboscis monkey populations as the ecological backbone of this species in fragmented landscapes. Throughout their distribution, proboscis monkeys are closely associated with riverine forests, which provide essential resources for feeding, sleeping, and daily movements (Bennett and Sebastian, 1988; Boonratana, 2013, 2000). Our spatial analyses further demonstrate that many of the remaining priority habitats, including mangrove forest, nipa palm forest, swamp forest, and lowland dry forest, occur immediately adjacent to existing protected areas, suggesting that relatively small-scale restoration or improved protection of these habitats could substantially enhance habitat continuity across the Lower Kinabatangan landscape. Notably, many of these suitable habitats have been maintained throughout the study period, indicating that their continued conservation should be prioritized through a combination of habitat protection and landscape-scale management. Such measures are likely to strengthen functional connectivity for proboscis monkeys while simultaneously benefiting a wide range of other floodplain-dependent species that persist within this highly fragmented ecosystem (Hearn et al., 2019).

Although our analyses focused on proboscis monkeys, the conservation priorities identified here are likely to benefit a much broader assemblage of floodplain wildlife. The Lower Kinabatangan supports one of the most important remaining populations of several threatened mammals, including Bornean orangutans (*Pongo pygmaeus*), Bornean elephants (*Elephas maximus borneensis*), and Sunda clouded leopards (*Neofelis diardi*), all of which are affected by habitat fragmentation and the loss of landscape connectivity (Ancrenaz et al., 2004; Goossens et al., 2016; Hearn et al., 2019). Consequently, restoring riparian forests and strategically maintaining suitable habitats beyond the existing protected-area network could generate conservation benefits well beyond a single flagship species. Because many of the priority restoration areas identified in this study occur adjacent to existing protected areas, relatively modest restoration efforts could substantially improve ecological connectivity across the Lower Kinabatangan floodplain (Goossens et al., 2016). Achieving these conservation outcomes will require conservation approaches that extend beyond protected areas and promote collaboration among government agencies, local communities, landowners, and other stakeholders across the wider Lower Kinabatangan landscape.

### 4.4 Strengths, limitations, and future directions

A major strength of this study is the integration of repeated landscape-wide population surveys, long-term monitoring at representative sites, and spatial analyses of habitat change throughout the study period. Such long-term, multidisciplinary datasets remain rare for tropical primates, despite growing recognition that reliable assessments of population trajectories and conservation effectiveness require sustained ecological monitoring through repeated observations (Barelli et al., 2023; Chapman et al., 2017, 2010). By combining independent lines of evidence, our study not only documents long-term population recovery but also identified the landscape-scale habitat changes associated with that recovery. This integrative approach provides a stronger basis for conservation planning than population or habitat assessments conducted in isolation.

Our study nevertheless has several limitations that also highlight important directions for future research. Although repeated boat-based surveys provide robust and repeatable estimates of population abundance and group structure, they do not allow direct estimation of demographic parameters such as survival, reproductive success, dispersal, or age-specific recruitment. Long-term studies of vertebrate populations have shown that integrating demographic processes with ecological monitoring is essential for understanding population persistence and conservation outcomes (Clutton-Brock and Sheldon, 2010). Likewise, although habitat change was quantified at the landscape scale, integrating long-term demographic, genetic, movement, and health data would provide a more comprehensive understanding of the mechanisms underlying population persistence. Our landscape-scale habitat analyses also do not account for finer-scale transition among habitat types, and changes within individual forest types may therefore be underrepresented. Collectively, these considerations highlight the value of multidisciplinary approaches, which are increasingly recognized as fundamental for advancing conservation science and adaptive management (Chapman and Peres, 2021).

More broadly, the framework presented here demonstrates how long-term ecological monitoring can be integrated with landscape-scale habitat assessment to evaluate conservation outcomes and identify practical priorities for adaptive management. As habitat fragmentation continues to threaten tropical biodiversity worldwide, similar integrative approaches will be essential for evaluating conservation outcomes, prioritizing restoration efforts, and guiding adaptive, evidence-based management in fragmented tropical landscapes. The Lower Kinabatangan demonstrates how sustained ecological monitoring can reveal conservation outcomes that would not be apparent from short-term studies alone.

## CRediT authorship contribution statement

**Muhammad Nur Fitri-Suhaimi:** conceptualization, data curation, funding acquisition, investigation, methodology, resources, visualization, formal analysis, writing – original draft. **Hiroshi Okamura**: methodology, writing – review and editing. **Nicola Karen Abram**: data curation, methodology, visualization, writing – review and editing. **Marc Ancrenaz**: data curation, resources, writing – review and editing. **Benoit Goossens**: resources, writing – review and editing. **Primus Lambut**: resources, writing – review and editing. **Joseph Tangah**: resources, writing – review and editing. **John C. M. Sha**: data curation, writing – review and editing. **Henry Bernard**: resources, writing – review and editing. **Vijay Subbiah Kumar**: resources, writing – review and editing. **Ikki Matsuda**: conceptualization, data curation, funding acquisition, investigation, methodology, project administration, resources, supervision, validation, writing – original draft.

## Declarations

### Ethics approval

The animal study was reviewed and approved by the Sabah Biodiversity Centre and the Sabah Wildlife Department [JKM/MBS.1000-2/2 JLD.20 (113)].

### Conflict of interest

The authors declare no competing interests.

### Generative AI and AI-assisted technologies in the writing process

During the preparation of this manuscript, the authors used DeepSeek (DeepSeek AI) and ChatGPT (OpenAI) to refine the clarity and logical flow of the text. The authors carefully reviewed, corrected, and approved all content generated, and take full responsibility for the final published version.

## Funding

This study was partially funded by Mandai Nature International Ltd. (no. GA2025041175 to M.N.F and I.M.), Core-to-Core Program, Asia-Africa Science Platforms (no. JPJSCCB20250006 to I.M.) and Japan Society for the Promotion of Science KAKENHI (no. 24H00774 to I.M.)

## Acknowledgement

The authors sincerely thank the Sabah Biodiversity Centre and the Sabah Wildlife Department for granting permission to conduct this research. We are deeply grateful to the Sabah Forestry Department for facilitating the use of its field facilities. Sonja Luz and Andie Ang (Mandai Nature International Ltd.) for their generous support in facilitating the research funding. We are grateful to Alexander Yee for providing accommodation and logistical support during the river surveys. We also sincerely thank Augustine Tuuga for his long-standing support and encouragement since the inception of this long-term research project. Finally, we thank our research assistants, Asnih Binti Etin and Jasrudy Bin Mandu, for their invaluable assistance during the fieldwork.

## Data availability

All data needed to evaluate the conclusions of this study are included in the article and its Supplementary Materials. Additional data are available from the corresponding author upon reasonable request.

## Notes

### Competing Interest Statement

The authors have declared no competing interest.

