## Supplementary Methods S1 for "Long-term Population Dynamics and Conservation of Proboscis Monkeys (*Nasalis larvatus*) in the Lower Kinabatangan, Sabah, Malaysia"

**Supplementary Methods S1. State-space model for long-term population dynamics**

Long-term population trends at the three monitoring sites were analysed using a state-space model that explicitly separates observation and process errors from the underlying population process (Buckland et al., 2004). Observed counts were assumed to follow a Poisson observation process,

$$Y_{t} \sim\mathrm{Poisson}\left( \mu_{t} \right)$$

where $Y_{t}$ is the observed number of proboscis monkeys at survey *t*, and

$$\mu_{t}=exp(x_{t})$$

where $x_{t}$ is the latent log-population state.

The latent log-population state was modelled as a first-order random walk,

$$x_{t} \sim N(a+ x_{t-1}, \sigma_{x}^{2})$$

where *a* represents the mean temporal trend in log population size and $\sigma_{x}^{2}$ is the process variance.

On the natural scale, this formulation is equivalent to exponential population dynamics:

$$N_{t+1}=\exp\left( a \right)\times N_{t}.$$

where exp (*a*) represents the multiplicative population growth rate between successive surveys.

Model parameters were estimated by maximum likelihood using Template Model Builder (Kristensen et al., 2016), which applies the Laplace approximation for efficient estimation of latent-variable models. Parameter estimation was performed separately for each of the three monitoring sites.

Predicted latent population trajectories were back-transformed to the natural scale and compared with the observed counts for visualization of long-term population trends. All statistical analyses were conducted in R version 4.4.1.
